# Modification and *de novo* design of non-ribosomal peptide synthetases (NRPS) using specific assembly points within condensation domains

**DOI:** 10.1101/354670

**Authors:** Kenan A. J. Bozhüyük, Annabell Linck, Andreas Tietze, Frank Wesche, Sarah Nowak, Florian Fleischhacker, Helge B. Bode

## Abstract

Many important natural products are produced by non-ribosomal peptide synthetases (NRPSs) ^1^.These giant enzyme machines activate amino acids in an assembly line fashion in which a set of catalytically active domains is responsible for the section, activation, covalent binding and connection of a specific amino acid to the growing peptide chain ^1,2^. Since NRPS are not restricted to the incorporation of the 20 proteinogenic amino acids, their efficient manipulation would give access to a diverse range of peptides available biotechnologically. Here we describe a new fusion point inside condensation (C) domains of NRPSs that enables the efficient production of peptides, even containing non-natural amino acids, in yields higher than 280 mg/L. The technology called eXchange Unit 2.0 (XU_2.0_) also allows the generation of targeted peptide libraries and therefore might be suitable for the future identification of bioactive peptide derivatives for pharmaceutical and other applications.

## Introduction

Secondary metabolite derived drugs have become essential agents to cure infectious diseases during the last almost 70 years^3,4^. Yet, infectious diseases are still the second major cause of death worldwide and furthermore, the world is facing a global public-health crisis as there is a growing risk of re-entering a pre-antibiotic era, since more and more infections are caused by multi-drug-resistant bacteria ^5^.

One source of new antibacterial agents are non-ribosomally made peptides (NRPs). Their high structural diversity imparts to them many properties of biological relevance and peptides have been identified with antibiotic, antiviral, anti-cancer, anti-inflammatory, immunosuppressant and surfactant qualities^6, 7, 8^. However, natural products often need to be modified to improve clinical properties and/or bypass resistance mechanisms^9,10^. To date, most clinically used NP derivatives are created by means of semi-synthesis^9,11^. A promising alternative strategy is the use of engineering approaches to modify NRP producing non-ribosomal peptide synthetase (NRPS) directly in order to produce optimized or non-natural natural products ^12^. However, to date most attempts to achieve this have yielded impaired or non-functional biosynthetic machineries ^7,13^.

NRPSs are large multienzyme complexes (megasynthases) ^14^that form peptides not limited to the twenty proteinogenic amino acids (AA)^15^. Furthermore, these NRPS can generate linear or cyclic peptides containing D-AA, N-methylated AA, N-terminal attached fatty acids (FA) or heterocycles^1,2,14,15^. NRPS do this by exhibiting a strict modular architecture in which a module is defined as the catalytic unit responsible for the incorporation of one specific building block (e.g. AA) into the growing peptide chain (N → C) and associated functional group modifications^16^. Modules are composed of domains that catalyze the single reaction steps like activation, covalent binding, optional modification of the building blocks, and condensation with the amino acyl or peptidyl group on the neighboring module^17^. At least three domains or essential enzymatic activities, respectively, are necessary for the non-ribosomal production of peptides (Fig. 1). They reside in the adenylation (A) for AA activation, thiolation (T) for AA tethering, and condensation (C) domains for peptide bond formation. Finally, most NRPS termination modules harbor a TE domain that releases the peptide, often in a cyclized form. These standard domains are additionally joined by tailoring domains that can catalyze epimerization (E), methylation (MT), cyclization (CY) or other modifications of the building blocks or the growing peptide chain^1^.

**Figure 1.**
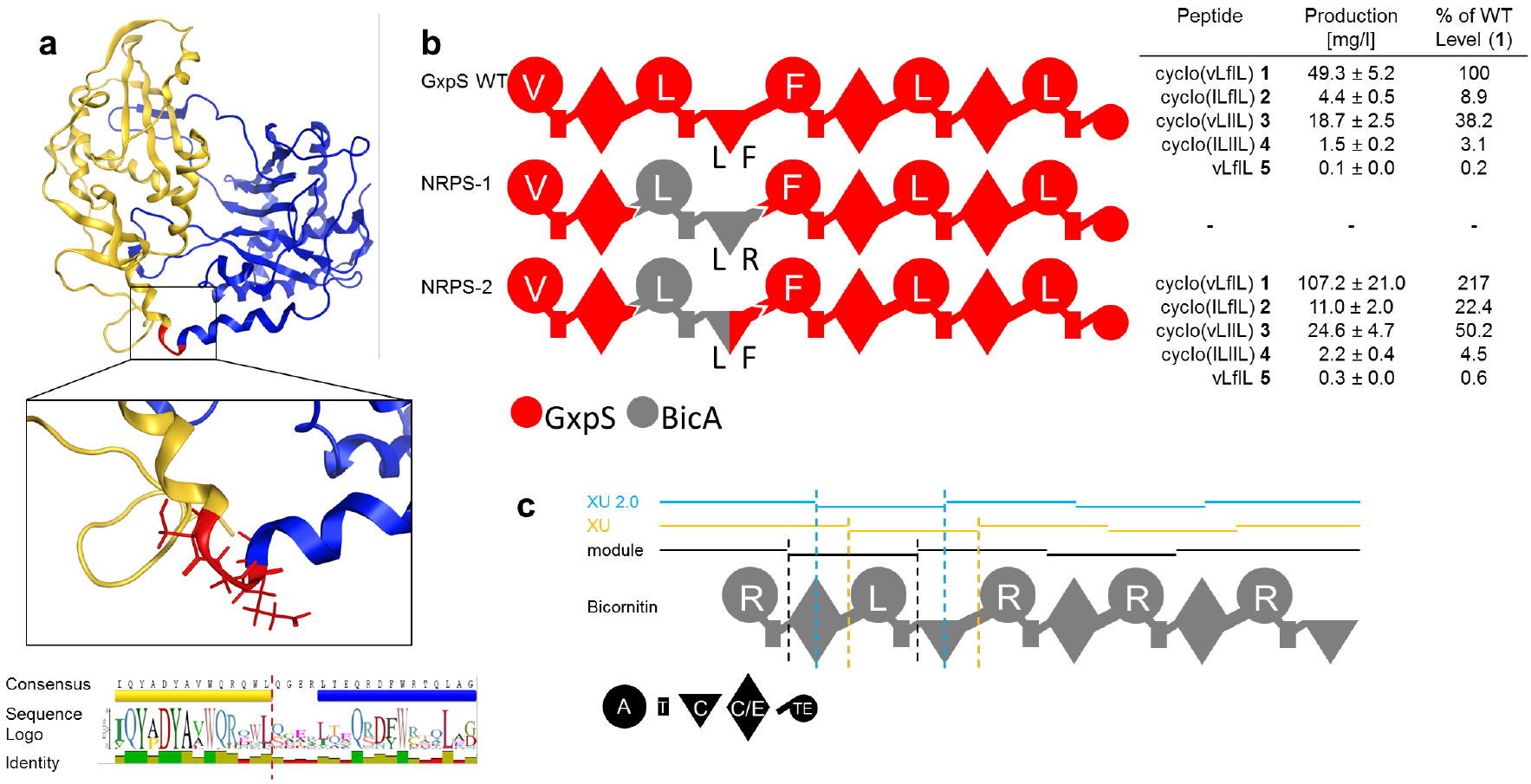
Modulation of C-domain substrate specificity. (**a**) C-domain excised from the T-C bidomain TycC 5-6 from tyrocidine syntethase (TycC) of *Brevibacillus brevis* (PDB-ID: 2JGP) with N-terminal (yellow) and C-terminal (blue) subdomains depicted in ribbon representation (top). Boxed: enlarged representation of the C_Dsub_ – C_Asub_ linker with contributing linker AAs in stick representation and fusion site marked in red. Bottom: sequence logo of C_Dsub_ – C_Asub_ linker sequences from *Photorhabdus* and *Xenorhabdus*. (**b**) Schematic representation of WT GxpS, recombinant NRPS-1 and −2 as well as corresponding peptide yields as obtained from triplicate experiments. For peptide nomenclature the standard one letter AA code with lowercase for D-AA is used. (**c**) Schematic representation of BicA with modules and eXchange Units (XU and XU_2.0_) highlighted. Specificities are assigned for all A-domains. For domain assignment the following symbols are used: A (large circles), T (rectangle), C (triangle), C/E (diamond), TE (C-terminal small circle).

Due to the modular character of the NRPS scientists strived to reprogram these systems via (I) the substitution of the A or paired A-T domain activating an alternative substrate, (II) the targeted alteration of just the substrate binding pocket of the A domain or (III) substitutions that treat C-A or C-A-T domain units as inseparable pairs ^7^. These strategies are complemented by recombination studies which have sought to re-engineer NRPS by T^18^, T-C-A^19^, communication domain^20^and A-T-C swapping^21^. However, with exception of the latter and recently published strategy, denoted as the concept of e<u>X</u>change <u>U</u>nits (XU)^22^, it has been difficult to develop clearly defined, reproducible and validated guidelines for engineering modified NRPS.

The limitation of the XU-concept is that the natural downstream C domain specificity must be obeyed clearly restricting its applicability and the C-domain specificities have to be met - at the donor as well as at the acceptor site. This disadvantage can be accepted if a large number of XUs with different downstream C domains are available. Due to these limitations also at least two XUs have to be exchanged to produce a new peptide derivative that differs in one AA position from the primary sequence of the wild type (WT) peptide^22^. However, a more flexible system reducing the limitations of C-domain specificities would drastically reduce the amount of NRPS building blocks necessary to produce or alter particular peptides and would enable the creation of artificial natural product libraries with hundreds or thousands of entities for large scale bioactivity screenings.

## Results and Discussion

### C-domains have acceptor site substrate specificity

To verify the influence of the C-domains acceptor site (C_Asub_) proof reading activity, the GameXPeptide producing NRPS GxpS of *Photorhabdus luminescens* TT01 (Supplementary Figure 1 and 2) was chosen as a model system ^23,24^. A recombinant GxpS was constructed, not complying with the C-domain specificity rules of the XU concept^22^. Here, XU2 of GxpS (Fig. 1b, **NRPS-1**) was exchanged against XU2 of the bicornutin producing NRPS (BicA, Fig. 1c) ^25^. Although both XUs are Leu specific, they are differentiated by their C_Asub_ specificities - Phe for XU2 of GxpS and Arg for XU2 of BicA. Therefore, no peptide production was observed as expected. This experiment confirmed previously published scientific results from *in vitro* experiments^26–29^, and illustrates that C domains indeed are highly substrate specific at their C_Asub_. From the available structural data of C domains it is clear that they show a pseudo-dimer configuration ^28,30–32^with their catalytic center, including the HHXXXDG motif, having two binding sites - one for the electrophilic donor substrate and one for the nucleophilic acceptor substrate ^29^(Fig. 1a and Supplementary Figure 3). Therefore we concluded that the four AA long conformationally flexible loop/linker between both subdomains might be the ideal target to reconfigure C domain specificities via the engineering of C domain hybrids (Fig. 1a). For this purpose the Arg specific C_Asub_ of the GxpS-BicA hybrid NRPS (Fig. 1b, **NRPS-1**) was re-exchanged to the Leu specific C_Asub_ of GxpS, restoring the functionality of the hybrid NRPS (**NRPS-2**) and leading to the production of GameXPeptide A-D (**1**-**5**) in 217% (107 mg/L) yield compared to the WT GxpS (Fig. 1b) as confirmed by MS/MS analysis and comparison of the retention times with a synthetic standard (Supplementary Figure 4).

**Figure 2.**
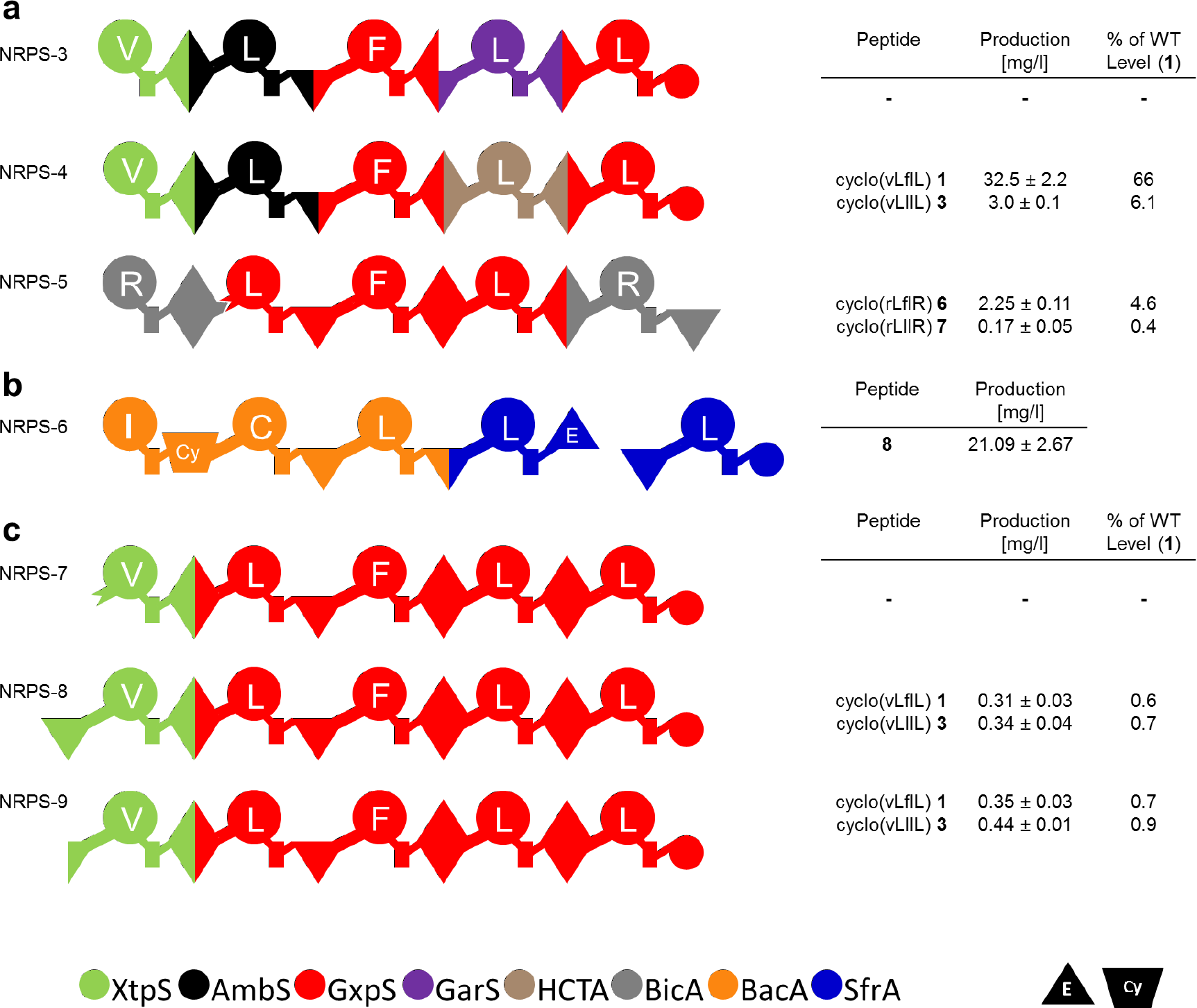
Design of recombinant NRPS for peptide production. (**a**) Generated recombinant GxpS (**NRPS-3** - **-5**) and corresponding amounts of GameXPeptide derivatives **1**, **3**, **6**, and **7** as determined in triplicates. (**b**) Recombinant NRPS-6 synthesizing **8**. Building blocks are of Gram-positive origin. (**c**) Schematic representation of recombinant GxpS (**NRPS-7** - **-9**) and corresponding peptide yields as obtained from triplicate experiments. For peptide nomenclature the standard AA one letter code with lowercase for D-AA is used. For assignment of domain symbols see Fig. 1; further symbols are E (epimerization; inverted triangle), CY (heterocyclization; trapezium). Bottom: Color code of NRPS used as building blocks (for details see Supplementary Figure 5).

**Figure 3.**
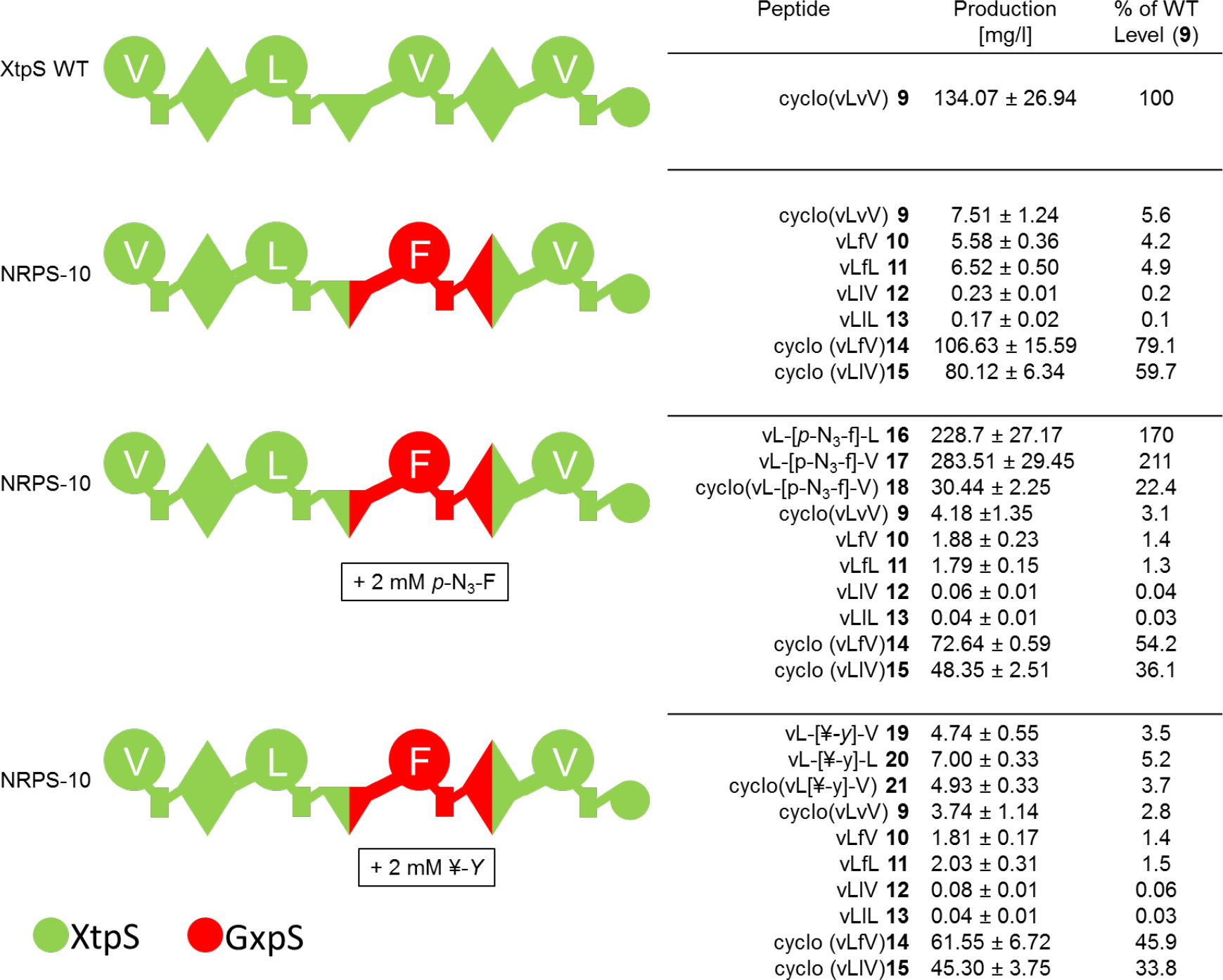
Creation of functionalized xenotetrapeptide derivatives. Schematic representation of WT XtpS, recombinant **NRPS-10** and corresponding peptide yields as obtained from triplicate experiments. For peptide nomenclature the standard one letter AA code with lowercase for D-AA is used. For assignment of domain symbols see Fig. 1. Bottom: Color code of NRPS used as building blocks (for details see Supplementary Figure 5).

**Figure 4.**
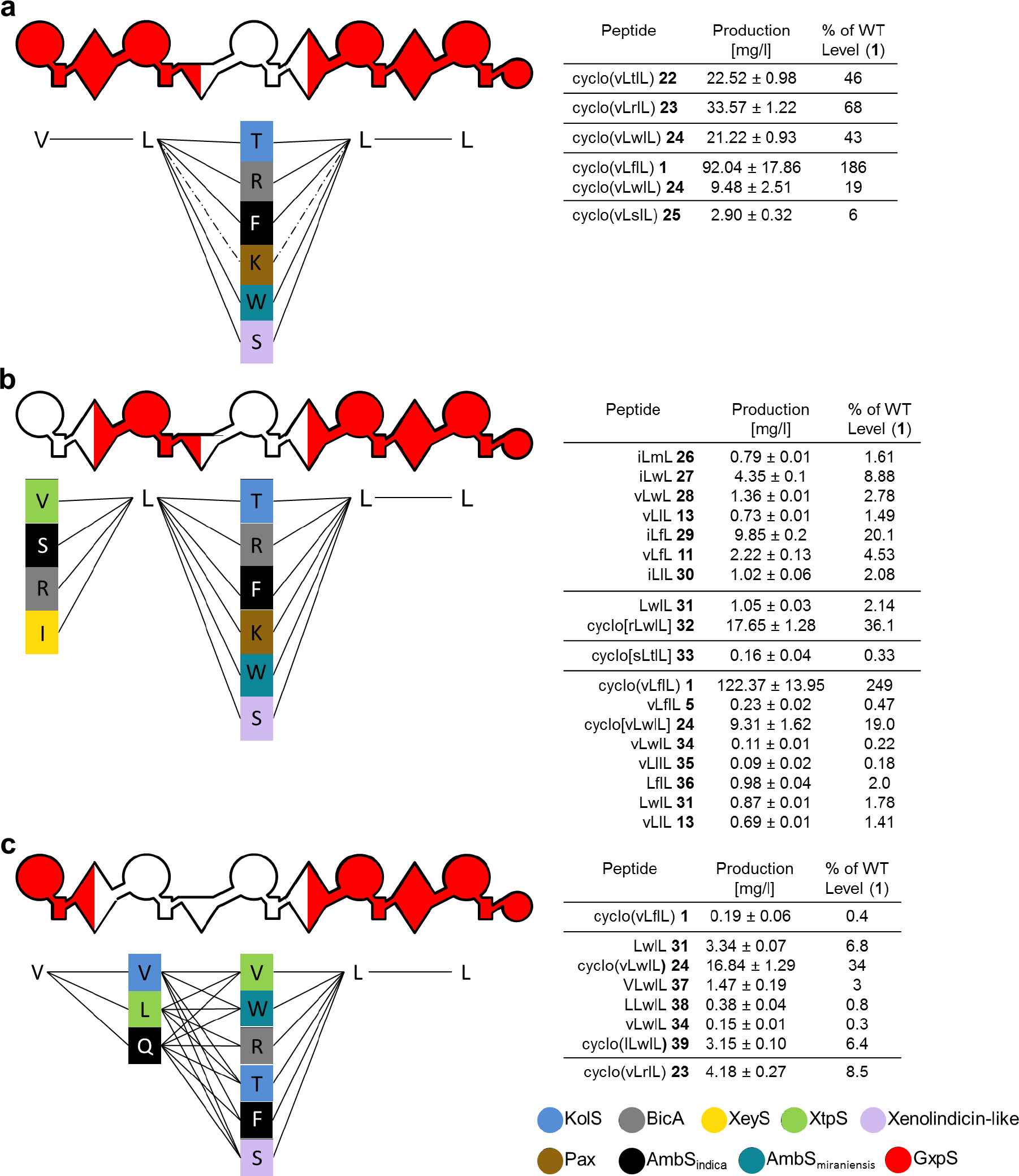
Targeted randomization of GxpS. Schematic representation of all possible recombinant NRPSs and corresponding NRPs (left). Detected peptides and corresponding peptide yields (right) as obtained from triplicate experiments. For peptide nomenclature the standard one letter AA code with lowercase for D-AA is used. For assignment of domain symbols see Fig. 1. Bottom: Color code of NRPS used as building blocks (for details see Supplementary Figure 5). (**a**) Randomization of position three from GxpS. (**b**) Randomization of position one and three from GxpS. (**c**) Randomization of adjacent positions two and three.

### The e*X*change *U*nit *2.0* concept

From these results in conjunction with bioinformatics analysis, we concluded that C-domains acceptor and donor site (C_Dsub_) mark a self-contained catalytically active unit C_Asub_-A-T-C_Dsub_ (XU_2.0_) without interfering major domain-domain interfaces/interactions during the NRPS catalytic cycle ^33^. In order to validate the proposed XU_2.0_ building block (Fig. 1c) and to compare the production titers with a natural NRPS, we reconstructed GxpS (Fig. 1b) in two variants (Fig. 2a, **NRPS-3** and **-4**). Each from five XU_2.0_ building blocks from four different NRPSs (XtpS, AmbS, GxpS, GarS, HCTA) (Supplementary Figure 5):

**NRPS-3** showed a mixed C/E_Dsub_-C_Asub_-domain between XU_2.0_3 and XU_2.0_4 (Fig. 2a), to reveal if C and C/E domains can be combined. In **NRPS-4** XU_2.0_3 from HCTA instead of GarS was used in order to prevent any incompatibilities (Fig. 2a).

Whereas **NRPS-3** (Fig. 2a) showed no detectable production of any peptide, **NRPS-4** (Fig. 2) resulted in the production of **1** and **3** in 66 and 6 % yield, respectively, compared to the natural GxpS, as confirmed by MS/MS analysis and comparison of the retention times of synthetic standards (Fig. 2a, Supplementary Figure 6). In line with expectations from domain sequences, phylogenetics, as well as structural idiosyncrasies of C/E- and C-domains ^29^, it may be deduced from these results that C/E and C-domains cannot be combined with each other. Although **NRPS-4** (Fig. 2a) showed moderately reduced production titers, most likely due to the non-natural C_Dsub_-C_Asub_ pseudo-dimer interface, the formal exchange of the promiscous XU_2.0_1 from GxpS (for Val/Leu) against the Val-specific XU_2.0_1 from XtpS led to exclusive production of **1** and **3** (Fig. 2a) without production of **2** and **4** observed in the original GxpS (Fig. 1b), indicating that the XU_2.0_ can also be used to increase product specificity and to reduce the formation of side products.

Additional GameXPeptide derivatives were generated (Fig. 2a, **NRPS-5**) by combining building blocks according to the definition of XU^22^ and XU_2.0_. Three fragments (1: C1-A1-T1-C/E2 of BicA; 2: A2-T2-C3-A3-T3-C/E4-A4-T4-C/E_Dsub_5 of GxpS; 3: C/E_Asub_5-A5-T5-C_term_ of BicA) from two NRPSs (BicA: *Xenorhabdus budapestensis* DSM 16342; GxpS: *Photorhabdus luminescens* TT01) were used as building blocks ^23,25^. The expected two Arg containing cyclic pentapeptides **6** and **7** were produced in yields of 4.6 and 0.4 mg/L and were structurally confirmed by chemical synthesis (Supplementary Figure 7). Both peptides only differ in Leu or Phe at position three from the promiscous XU_2.0_3 from GxpS. Despite a drop of production rate in comparison to the WT NRPS, we successfully demonstrated that the recently published XU^22^ as well as the novel XU_2.0_ strategy can be combined for successful reprogramming of NRPS and the production of tailor-made peptides.

To show the general applicability of the novel XU_2.0_ building block an artificial NRPS was designed *de novo* from building blocks of Gram-positive origin (NRPS for the production of bacitracin^34^from *Bacillus licheniformis* ATCC 10716 and surfactin^35^from *Bacillus subtilis* MR 168), since all aforementioned recombined NRPS are of Gram-negative origin. The expected pentapeptide **8** containing the bacitracin NRPS derived thiazoline ring was produced in yields of 21.09 mg/L (Fig. 2b, Supplementary Figure 8) in *E. coli*, showing the universal nature of the XU_2.0_.

### Amending the starter unit

Up to date there is no publication describing the successful exchange of a starter unit against an internal NRPS-fragment. Reasons for that might be that (I) starter-A-domains in general comprise some kind of upstream sequence of variable length with unknown function and structure, which makes it difficult to define an appropriate artificial leader sequence, and that (II) necessary interactions at the C-A interface may be important for adenylation activity and A-domain stability as indicated recently^36,37^. In order to test whether the XU_2.0_ concept can also be applied to modify starter units, three recombinant GxpS constructs (**NRPS-7** - **9**) with internal domains as starting units were created (Fig. 2c). In **NRPS-7** A1-T1-C_Dsub_2 of GxpS was exchanged against C2A3-linker-A3-T3-C_Dsub_4 of XtpS since all starter A-domains have at least a preceding C-A linker sequence. Since there are several examples of NRPSs carrying catalytically inactive starter C-domains (e.g. AmbS) ^38^, A1-T1-C_Dsub_2 of GxpS was altered to C3-A3-T3-C_Dsub_4 of XtpS in **NRPS-8**. In **NRPS-9** A1-T1-C_Dsub_2 of GxpS was altered to C_Asub_-A3-T3-C_Dsub_4 of Xtps since there are natural NRPSs exhibiting parts of a C-domain (e.g. BicA) in front of the starter A-domain.

Whereas **NRPS-7** (Fig. 2c) did not show production of the desired peptides, **NRPS-8** and **NRPS-9** synthesized **1** and **3** in yields between 0.31-0.44 mg/L (Fig. 2c, Supplementary Figure 9). This indicates that internal A-domains can indeed be used as starter domains, if the upstream C_Asub_ or C-domain is kept in front of the A-domain pointing to the importance of a functional C-A interface for A-domain activity. Yet, the observed low production titers might indicate that for example the observed difference in codon usage and the lower GC-content at the beginning of WT NRPS encoding genes could have a major impact on transcriptional and/or translational efficiency in conjunction with protein folding as described previously^39,40^.

### Production of functionalized peptides

Besides simply creating NRP derivatives, one useful application of NRPS reprogramming is the incorporation of non-proteinogenic or even non-natural AA. Examples for the latter might be AAs containing alkyne or azide groups, allowing reactions like Cu(I)-catalyzed or strain-promoted Huisgen cyclization also known as “click” reactions ^41,42,42–44^. Yet, although NPRS and A domains have been examined exhaustively for several years, no general method for the *in vivo* functionalization of NRPs are available by reprogramming NRPS templates.

A broad range of AAs are accepted by the A3 domain of GxpS (Supplementary Figure 10) resulting in a large diversity of natural GameXPeptides ^23,24^. Moreover, by using a ɣ-^18^O_4_-ATP pyrophosphate exchange assay for A domain activity^45,46^ and adding substituted phenylalanine derivatives to *E. coli* cultures expressing GxpS, the respective A3-domain was identified as being able to activate (*in vitro*, Supplementary Figure 10) and incorporate (*in vivo*, Supplementary Figure 11) several *ortho*- (*o*), *meta*- (*m*) and *para*- (*p*) substituted phenylalanine derivatives, including 4-azido-L-phenylalanine (*p*N_3_-F) and *O*-propargyl-L-tyrosine (¥-Y). When the Val specific XU_2.0_3 of the xenotetrapeptide^47^ (**9**) (Supplementary Figure 12) producing NRPS (XtpS) from *X. nematophila* HGB081 was exchanged against XU_2.0_3 of GxpS, six new xenotetrapeptide derivatives (**10**-**15**) in yields between 0.17-106 mg/L were produced reflecting its natural promiscuity (Fig. 3, Supplementary Figure 13). After adding *p*N_3_-F and ¥-Y to growing *E. coli* cultures expressing recombined XtpS (**NRPS-10**), six functionalized peptides (**16**-**21**) differing in position 3 were produced in yields of 5-228 mg/L with **17**, **18** and **19** being structurally confirmed by ch
emical synthesis. Moreover, although the A4-domain of XtpS shows an exclusive specificity for Val in the WT NRPS XtpS, peptides **11**, **13**, **16** and **21** produced by **NRPS-10** additionally incorporate Leu at position four. The observed change in specificity might be due to the hybrid C/E4-domain upstream of A4 from **NRPS-10**. Leu is the original substrate downstream of the introduced XU_2.0_3 of GxpS (Fig. 1b) in its natural context, indicating that the overall structure of C domains along with resulting transformed C-A interface interactions might influence the A domain substrate specificity. Recently, similar but *in vitro* observed effects were reported regarding A domains from sulfazecin^36^and microcystin^37^. This effect could also be used to increase the specificity of A domains to prevent the formation of side products. Further investigations will shed light on this remarkable and yet unreported effect.

### Production of peptide libraries

Modern drug-discovery approaches often apply the screening of compound libraries including NP libraries^48^since they exhibit a wide range of pharmacophores, structural diversity and have the property of metabolite-likeness often providing a high degree of bioavailability. Yet, the NP discovery process is as expensive as time consuming^49^. Consequently, for bioactivity screenings the random recombination of certain NRPS fragments would be a powerful tool to create focused artificial NP-like libraries.

In an initial test, GxpS was chosen for the generation of a focused peptide library created via a one-shot yeast based TAR cloning approach^38,50^. Here, the third position of the peptide (D-Phe) was randomized (Fig. 4a) using six unique XU_2.0_ building blocks from six NRPS (KolS^51^, AmbS_mir_^38^, Pax^52^, AmbS_ind_ XllS; for detail see Supplementary Figure 5), resulting in the production of **1** and four new GameXPeptide derivatives (**22-25**) in yields of 3-92 mg/L that were structurally confirmed by chemical synthesis (Supplementary Figure 14).

For the generation of a second and structurally more diverse peptide library, positions 1 (D-Val) and 3 (D-Phe) of GxpS were selected in parallel for randomization (Fig. 4b). From the experimental setup theoretically 48 different cyclic or linear peptides could be expected. Screening of 50 *E. coli* clones resulted in the identification of 18 unique cyclic and linear peptides (**1**, **5**, **11**, **13**, **24**, **26**-**36**) from four peptide producing clones differing in peptide length and AA composition (Supplementary Figure 15). Since only 7 from 18 identified peptides belong to the originally expected set of peptides, homologues recombination in yeast based reprogramming of NRPS also allows the production of unexpected peptides due to unexpected homologues recombination events resulting in an additional layer of peptide diversification, as observed previously ^22^.

Randomizing directly adjacent positions via a similar approach requires a standardized nucleotide sequence (40 base pairs) for homologues recombination (Supplementary Figure 16) ^38,50^. From a detailed analysis of the T-C didomain crystal structure of TycC5-6 (PDB-ID: 2JGP), helix α5 (I253-F265) next to the C domain’s pseudo-dimer linker was identified as an ideal target for homologues recombination. Subsequently, an artificial α5 helix was designed to randomize position 2 (L-Leu) and 3 (D-Phe) of GxpS (Supplementary Figure 16a), being an integral part of all resulting recombinant C3 domains and therefore connecting XU_2.0_2 and 3. The applied α5 helix was defined as the consensus sequence of all involved XU_2.0_ building blocks (Supplementary Figure 16b). Screening of 25 *E. coli* clones revealed the synthesis of eight cyclic and linear GameXPeptides (**1**, **23**-**24**, **31**, **34**, **37**-**39**) from three peptide producing clones in good yields, showing the general applicability of redesigning α5 with respect to randomly reprogramming biosynthetic templates (Fig. 4c and Supplementary Figure 17).

### Conclusion

We have recently published the XU concept enabling the efficient reprogramming of NRPS but limited in its applicability by downstream C domain specificities. Here we present the XU_2.0_ concept that eliminates these limitations by a direct assembly inside the C domains and allows the production of natural and artificial peptides in yields up to 280 mg/L. For the construction of any peptide based on the 20 proteinogenic AAs only 80 XU_2.0_ building blocks are necessary (only four of each: C_Dsub_-A-T-C_Asub_, C_Dsub_-A-T-C/E_Asub_, C/E_Dsub_-A-T-C/E_Asub_, and C/E_Dsub_-A-T-C_Asub_) whereas 800 building blocks would be necessary to generate the same number of peptides using the XU concept. Consequently, the introduction of the XU_2.0_ simplifies and broadens the possibilities of biotechnological applications with respect to optimize bioactive agents via NRPS engineering exemplified for the reprogramming of NRPSs (Fig. 1 and 2) or the production of functionalized peptides by incorporating XU_2.0_ building blocks accepting non-natural AAs like *p*N_3_-F and ¥-Y (Fig. 3, Supplementary Figure 9) allowing further derivatization^41,42,53^.

However, the true strength of the XU_2.0_ concept is its application to generate random NP-like peptide libraries (Fig. 4) for subsequent bioactivity screenings. The possible automation of NRPS library design coupled to a bioactivity screening opens up entirely new opportunities of identifying novel lead compounds in the future. Especially in the area of anti-infective research the XU_2.0_ concept might allow a fast access to natural product derivatives with altered bioactivity profiles or for the generation of producer strains with less side products to facilitate compound purification.

**Supplementary Information** is available in the online version of the paper.

## Acknowledgements

The authors are grateful to Melanie Lindner and Caspar Zizka for help with the construction of selected constructs and Dr. Carsten Kegler for helpful discussions. This work was funded in part by a European research starting grant under grant agreement no. 311477 and LOEWE program of the state of Hesse as part of the MegaSyn research cluster. H.B.B. acknowledges the Deutsche Forschungsgemeinschaft for funding of the Impact II qTof mass spectrometer (INST 161/810-1).

## Author Contributions

K.A.J.B. and H.B.B. designed the experiments. K.A.J.B., A.L., A.T. and S.N. performed all molecular biology and biochemical experiments, F.W. synthesized all peptide standards that were used for HPLC/MS-based quantification performed by A.L. All authors analysed the results and K.A.J.B., A.L. and H.B.B. wrote the manuscript. All authors saw and approved the manuscript.

